# Atrial Granules in Atrial Cardiomyocytes as Acidic Calcium Stores

**DOI:** 10.1101/2024.11.25.625237

**Authors:** Emily Akerman, Daniel Aston, Eva Rog-Zielinska, Barry Boland, Ulrich Schotten, Sander Verheule, Rebecca A Capel, Rebecca A.B. Burton

## Abstract

Acidic calcium stores significantly influence basal calcium transient amplitude and β-adrenergic responses in cardiomyocytes. Atrial myocytes express a small acidic organelle called atrial granules (AG), which store and excrete atrial natriuretic peptide and are not expressed by healthy ventricular myocytes. AG are known to be acidic with a high calcium content. The number and position of these calcium-rich organelles relative to other signaling sites has not been investigated. Staining of acidic organelles in adult guinea pig cardiomyocytes showed the presence of fluorescent acidic puncta throughout the cytosol. Atrial myocytes exhibited an increased concentration of acidic organelles at the nuclear poles. Live cell fluorescent studies using PBA to inhibit peptidylglycine α-amidating monooxygenase, a crucial component of AG membranes, effectively eliminated staining at the nuclear poles and most acidic puncta in atrial cells. The application of PBA to ventricular myocytes did not affect LysoTracker staining. Electron microscopy studies on goat atrial fibrillation (AF) and sham control tissue, allowed visualization of AGs. Quantitative analysis revealed AGs to be in close apposition to the sarcoplasmic reticulum and mitochondria. AGs were significantly increased in AF goat samples when compared to sinus rhythm from 3D electron tomography images. Our imaging studies suggest that AGs make up a large percentage of atrial acidic stores, with AG associated with the sarcoplasmic reticulum and their number increasing during AF. We raise the question whether the positioning of AGs are strategic to communicate with other calcium organelles. Further studies to investigate whether AGs contribute to physiological calcium signalling are required.

## Introduction

In addition to their degradative role in cells, acidic organelles (namely lysosomes) contribute significantly to basal calcium transient amplitude and β-adrenergic responses in both atrial [1] and ventricular myocytes [2]. Atrial granules (AGs) are also known to be acidic and contain a high calcium content [3]. Lysosomes, in ventricular myocytes, are located near the sarcoplasmic reticulum (SR) and mitochondria, creating membrane contact sites that can potentially function as signaling microdomains [4]. The discovery of AGs and natriuretic peptides within them lead to the atrium being identified as an endocrine organ [5]. AGs are lysosome-related organelles which secrete atrial natriuretic peptide (ANP). Peptylglycine α-amidating monooxygenase (PAM) [6] is the primary membrane protein in atrial granules [6], crucial for amidated peptide biosynthesis [6]. PAM plays a noncatalytic role in the atrium and its actions are confined to the early stages of the secretory pathway [7]. An accumulation of AGs around the golgi complex has been observed [3], however their number relative to lysosomes and positioning with relation to other calcium signalling sites has not been fully explored in atrial physiology and pathophysiology.

## Methods

### Myocyte Isolation

All animal experiments were performed in accordance with the United Kingdom Home Office Guide on the Operation of Animal (Scientific Procedures) Act of 1986. Live cell imaging experiments were conducted on Guinea pig isolated adult atrial and ventricular cardiomyocytes. Cell isolation methods were previously published in [2]. Briefly, guinea pig hearts were dissected and washed in heparin-containing PSS (in mM: NaCl 125, NaHCO_3_ 25, KCl 5.4, NaH_2_PO_4_ 1.2, MgCl_2_ 1, glucose 5.5, CaCl_2_ 1.8, pH to 7.4 with NaOH, 20 IU heparin per mL to prevent blood clots forming) and mounted on a Langendorff setup for retrograde perfusion via the aorta. The heart was perfused in a modified Tyrode solution containing (in mM): NaCl 136, KCl 5.4, NaHCO_3_ 12, sodium pyruvate 1, NaH_2_PO_4_ 1, MgCl_2_ 1, EGTA 0.04, glucose 5; gassed with 95% O_2_/5% CO_2_ to maintain a pH of 7.4 at 37°C for 3 minutes. Solution was switched to a digestion solution: the modified Tyrode above containing 100 µM CaCl_2_ and 0.04 mg/ml Liberase™ (Roche, Penzberg, Germany) and no EGTA. After 25 minutes of enzymatic digestion, the heart was removed. Atria were separated from ventricles in a dissection bath. Tissue was cut into small pieces (around 2×2 mm^3^) followed by 1 minute of gentle trituration. Atrial myocytes were stored in a high potassium medium (in mM: KCl 70, MgCl_2_ 5, K^+^ glutamine 5, taurine 20, EGTA 0.04, succinic acid 5, KH_2_PO_4_ 20, HEPES 5, glucose 10; pH to 7.2 with KOH0) and ventricular myocytes were stored in modified Tyrode solution. Myocytes were left at 4°C to rest for 30 minutes before recordings and were used up to 4 hours post isolation.

### Live Cell Spinning Disc Confocal Studies

Lysotracker™ staining of acidic organelles (ThermoFisher, 100 nM, 20 minutes) in isolated guinea pig myocytes was performed to analyse acidic punctate organelles throughout the cytoplasm. We used 4-Phenyl-3-butenoic acid (PBA, a turnover-dependent inactivator of PAM), to perturb AG formation (Sigma, 200 µM, 2h) in atrial and ventricular myocytes.

### High Resolution Three-Dimensional Electron Tomography

3D electron tomography imaging data on goat atrial tissue that was performed in previous studies [8] were used to analyse AGs. The goat model in this study (female C. hircus) is described in [9], AF was induced and maintained in female goats (C. hircus) for 6 months. Goats were anesthetized and euthanized following an open chest experiment (N=4 AF and N=4 sham controls). Atrial goat biopsies were either snap frozen or chemically fixed with Karnovsky fixative for electron microscopy studies. The goat study was carried out in accordance with the principles of the Basel declaration and regulations of European directive 2010/63/EU, and the local ethical board for animal experimentation of the Maastricht University approved the protocol.

### Statistics

Datasets were analysed using non-parametric Mann-Whitney U test. Datasets are presented as mean ± SEM. Differences were considered statistically significant at values of P<0.05, *P<0.05, **P<0.01, ***P<0.001, ****P<0.0001. Statistical analyses were performed using Prism 10 (GraphPad, CA, USA).

## Results

### Atrial Granules comprise the majority of acidic organelles in live atrial myocytes

Staining of acidic organelles by LysoTracker in freshly isolated guinea pig atrial and ventricular myocytes revealed fluorescent acidic puncta throughout the cytosol. In atrial myocytes there was an additional concentration of acidic organelles at the nuclear poles, consistent with published literature on the site of AG formation [7]. Inhibition of PAM (200 µM PBA), an essential component of AG membranes [7], abolished not only staining at the nuclear poles but the majority of acidic puncta in atrial myocytes (Figure 1A). In control conditions atrial myocytes had a perinuclear acidic puncta count of 22.7±1.8 per cell (n=10, Figure 1Bi), a peripheral count of 10.7±.2.3 (n=10, Figure 1Bii) at the top of the cell and 11.7±1.4 (n=10, Figure 1Biii) at the bottom of the cell. Addition of PBA caused a significant decrease in both perinuclear and peripheral cytosol; perinuclear acidic puncta count of 9.4±2.7 (n=14, ***P<0.001, Figure 1Bi), peripheral count of 3.5±1.7 (n=14, *P<0.05, Figure 1Bii) at the top and 6.2±1.9 (n=14, *P<0.05, Figure 1Biii) at the bottom. Application of PBA to ventricular myocytes had no impact on LysoTracker staining (Figure Figure 1C). In control conditions ventricular myocytes have a perinuclear acidic puncta count of 15.2±2.6 and a peripheral count of 17.5±1 1 at the top and 17.2±3.2 at the bottom of the cell (n=6, Figure 1Ci-iii). Addition of PBA had no significant change to the perinuclear or peripheral puncta count 19.8±2.9, 12.8±1 6 (top) and 13.7±1.6 (bottom) in ventricular myocytes (n=12, ns, Figure 1Ci-iii).

**Figure 1.**
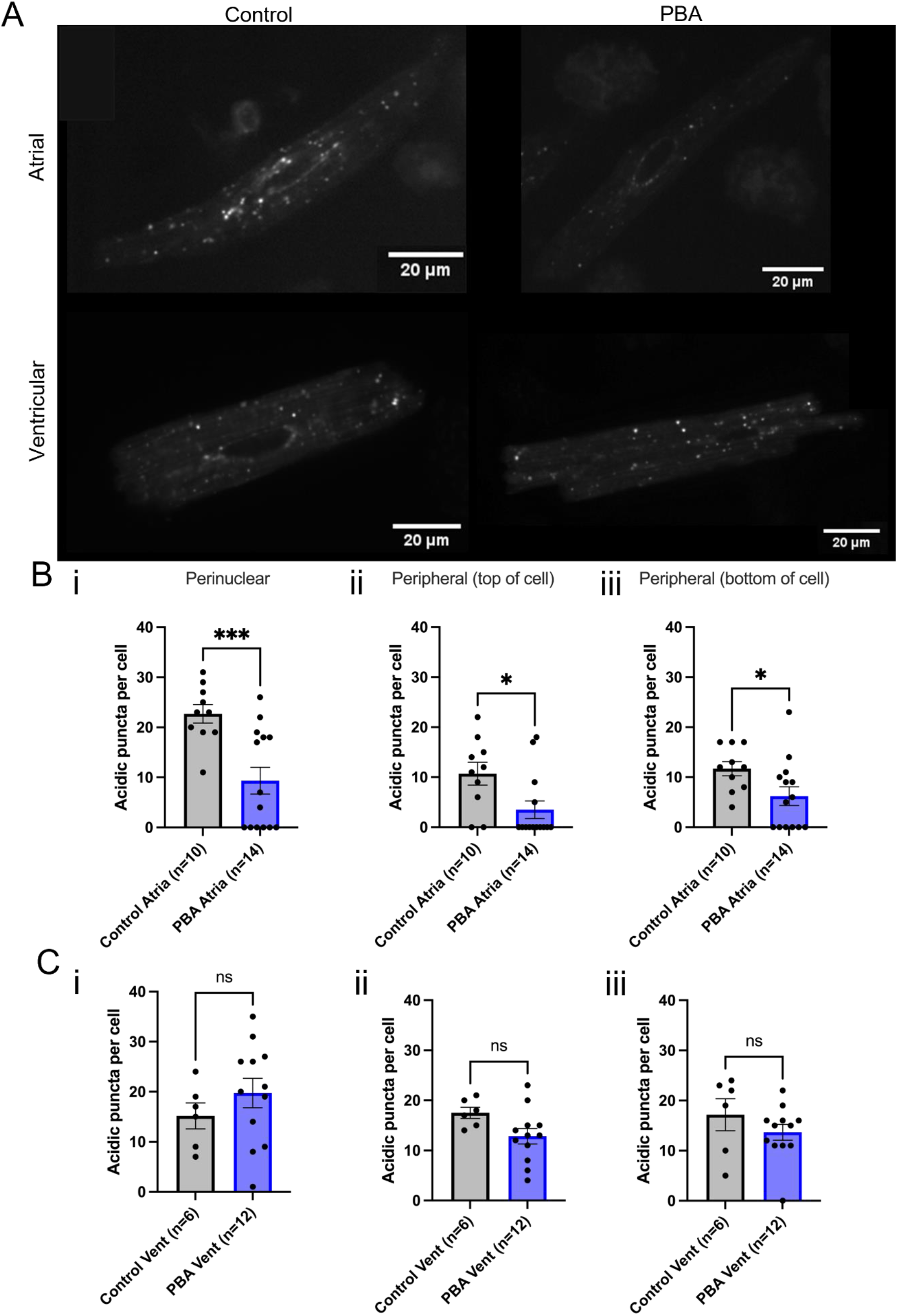
Atrial Granules comprise the majority of acidic organelles in live atrial myocytes. A: Lysotracker staining of acidic organelles (100 nM, 20 minutes) in primary isolated guinea pig myocytes throughout the sarcoplasm of atrial myocytes in control conditions (i) and in the presence of PBA (200 µM, 2h, ii), and in ventricular myocytes in control conditions (iii) and in the presence of PBA (200 µM, 2h, iv). Bi: Acidic perinuclear puncta count in control conditions (EtOH, n=10) and in the presence of PBA (200 µM, n=14) in atrial myocytes. ii: Acidic peripheral puncta count in control conditions (EtOH, n=10) and PBA (200 µM, n=14) at top of atrial myocytes. iii: Acidic peripheral puncta count in control conditions (EtOH, n=10) and PBA (200 µM, n=14) at bottom of atrial myocytes. Ci: Acidic perinuclear puncta count in control conditions (EtOH, n=6) and PBA (200 µM, n=12) in ventricle myocytes. ii: Acidic peripheral puncta count in control conditions (EtOH, n=6) and PBA (200 µM, n=12) at the top of ventricle myocytes. iii: Acidic peripheral puncta count in control conditions (EtOH, n=6) and PBA (200 µM, n=12) at the bottom of ventricle myocytes. PBA: 4-Phenyl-3-butenoic acid.

### Atrial Granule position changes significantly in a goat model of atrial fibrillation

We measured the size and position of AGs in relation to both the SR and mitochondria in left atrial tissue in a goat AF model using 3D electron tomography. Consistent with our live cell results, EM of fixed goat atrial tissue revealed the presence of both lysosomes and AG in atrial myocytes. AG in sham-operated control samples had a maximum diameter of 170±4 nm (n=35 lysosomes, N=4 goats, Figure 2A). AGs were observed in close apposition to the SR (13.8±1.72 nm at the closest point, n=34, N=4, Figure 2B). The average AG was a minimum of 195.6±34.8 nm (n=34, N=4) away from the nearest mitochondria (Figure 2C). After 6 months in AF, AGs showed no significant changes in diameter than those observed in sham-operated controls (maximum diameter 164±3 nm, n=50, N=4 goats, P>0.05, Figure 2A). In AF tissue, AGs were positioned significantly further away from their nearest SR, at an average distance of 25.1±2.61 nm (n=46, N=4), compared to 13.8±1.72 nm in sham-operated samples (***P<0.001, Figure 2B). AG in AF goats were significantly closer to mitochondria than those in control cells, with the average closest distance of 99.3±19 nm (n=48, N=4, **P<0.01, Figure 2C). A Summary table of the measure taken is presented in Figure 2D and representative snap shots of atrial granules (AG, red) to Mitochondria (Mito, blue) and Sarcoplasmic Reticulum (SR, yellow arrows) from goat left atrial sinus rhythm control (Figure 2E) and AF (Figure 2F) from 3D tomograms.

**Figure 2.**
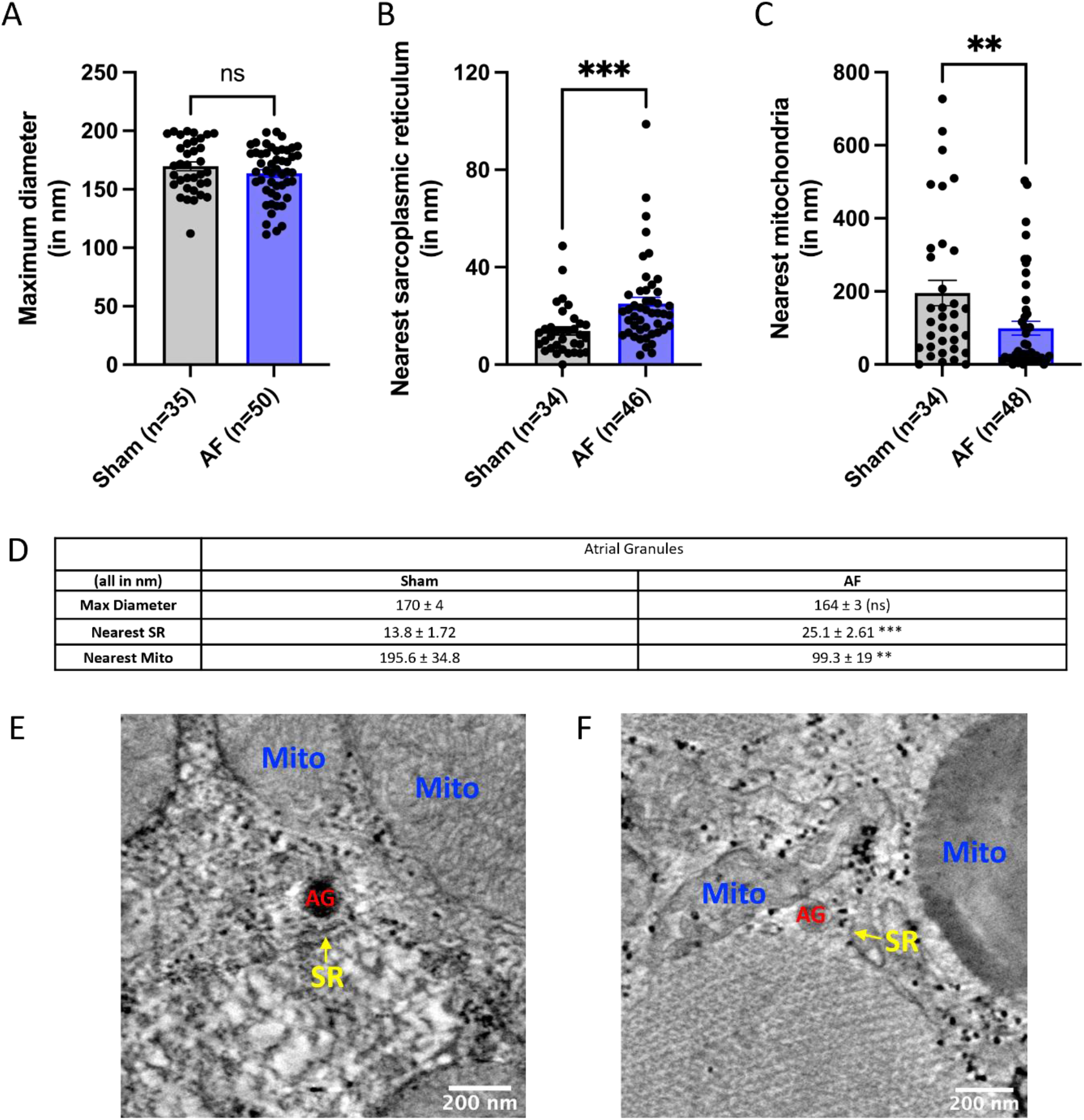
Atrial Granule position changes significantly in a goat model of atrial fibrillation. A: Atrial Granule (AG) size and positioning in goat AF tissue using 3D electron tomography. Maximum AG diameter in sham-operated (n=35) and AF goats (n=50). B: Minimum distance between AG and nearest SR measured in sham-operated (n=34) and AF goats (n=46). C: Minimum distance between lysosomes and mitochondria in sham-operated (n=34) and AF goats (n=48). D: Summary table indicating maximum diameter, nearest SR and nearest mitochondria measured from sham-operated and AF goats tomograms. AG: atrial granules; SR: sarcoplasmic reticulum. E: Snapshot from goat left atrial sinus rhythm control from 3D tomograms F: Snapshot from goat left atrial with AF from 3D tomograms. Snap shots from E and F show proximity of atrial granules (AG, red) to Mitochondria (Mito, blue) and Sarcoplasmic Reticulum (SR, yellow arrows). Scale bars provided on each image represent 400 nm.

## Discussion

Back in 1964, Jamieson and Palade [10] described AGs as “large populations (up to 600/cell) of spherical, electron-opaque granules approximately 0.3 to 0.4 micro in diameter are characteristically found in muscle fibers of mammalian atria”. Our imaging studies suggest that AGs comprise a large percentage of acidic calcium stores within atrial cardiomyocytes. Here for the first time, we describe nanojunctions between AGs and the SR in mammalian cardiomyocytes. In goat AF samples, we observe significant structural remodelling between the positioning of AGs to SR and mitochondria. We found that AGs are situated significantly further away from the SR and closer to mitochondria in goat AF tissue when compared to sham-operated controls (Figure 2). Similar observations have been made recently by our group in relation to positioning of lysosomes with mitochondria and SR [8]. The strategic positioning of AG raises the question of whether there is functional cross talk between AGs and other calcium-containing stores. It is an important next step to explore whether AGs contribute to physiological calcium signalling on top of their known hormone secretory role.

## Acknowledgements

R.A.B.B. is funded by a Sir Henry Dale Wellcome Trust and Royal Society Fellowship (109371/Z/15/Z) and R.A.B.B. acknowledges support from The Returning Carers’ Fund (Oxford University, Medical Sciences Division). R.A.B.B. is a Senior Research Fellow of at Linacre College. R.A.B.B. acknowledges research funds from the Ellis T Davies Fellowship Endowment, University of Liverpool. R.A.C. is funded by the Wellcome Trust and Royal Society, acknowledges the Returning Carer’s Fund and the Health Research Bridging Salary Scheme, University of Oxford. We thank Dr Errin Johnson and Raman Dhaliwal from the Dunn School of Pathology for Electron Microscopy technical support. The goat model work was supported by the Netherlands Heart Foundation (CVON2014-09, RACE V Reappraisal of Atrial Fibrillation: Interaction between hypercoagulability, Electrical remodeling, and Vascular Destabilisation in the Progression of AF, and Grant number 01-002-2022-0118, EmbRACE: Electro-Molecular Basis and the theRapeutic management of Atrial Cardiomyopathy, fibrillation and associated outcomEs), the European Union (CATCH ME: Characterizing Atrial fibrillation by Translating its Causes into Health Modifiers in the Elderly, grant number 633196; MAESTRIA: Machine Learning Artificial Intelligence Early Detection Stroke Atrial Fibrillation, grant number 965286). This research was funded in whole, or in part, by the Wellcome Trust [109371/Z/15/Z]. For the purpose of Open Access, the author has applied a CC BY public copyright licence to any Author Accepted Manuscript version arising from this submission.

## Author Contributions

R.A.B.B. and R.A.C. conceived and designed the study; S.V. and U.S. performed the goat AF experiments and provided the tissue for this study. R.A.B.B. and E.A. drafted the first draft of the paper. B.B and D.A. contributed intellectually to the project. E.A. and R.A.C. performed experiments. E.A. performed statistical analysis and prepared the figures. E.R-Z. and R.A.B.B. contributed to electron microscopy studies.

## Data availability

Data can be requested by contacting the lead contact. This study did not generate new unique codes.

## Inclusion & Ethics

This study follows inclusion and ethics in global research.

